# The precision of hippocampal representations predicts incremental value-learning across the adult lifespan

**DOI:** 10.1101/2025.04.08.647815

**Authors:** Camilla van Geen, Karolina M. Lempert, Michael S. Cohen, Kameron A. MacNear, Frances M. Reckers, Laura Zaneski, David A. Wolk, Joseph W. Kable

## Abstract

Correctly assigning value to different options and leveraging this information to guide choice is a cornerstone of adaptive decision-making. Reinforcement learning (RL) has provided a computational framework to study this process, and neural signals linked to RL have been identified in the striatum and medial prefrontal cortex. More recently, hippocampal contributions to this kind of value-learning have been proposed, at least under some conditions. Here, we test whether the hippocampus provides a signal of the option’s identity that aids in credit assignment when learning about several perceptually similar items, and evaluate how this process differs across the lifespan. A sample of 251 younger and older adults, including a subset (n = 76) with simultaneous fMRI, completed an RL task in which they learned the value of four houses through trial-and-error. Older adults showed decreased choice accuracy, accompanied by reduced neural signaling of value at choice but not feedback. Using representational similarity analysis, we found that the precision with which choice options were represented in the posterior hippocampus during choice predicted accurate decisions across age groups. Interestingly, despite previous evidence for neural de-differentiation in older adults, we found no support for a “blurring” of these stimulus representations in older adults. Rather, we observed reduced connectivity between the posterior hippocampus and the medial PFC in older adults, and this connectivity correlated with choice consistency. Taken together, these findings identify a hippocampal contribution to incremental value learning, and that reductions in incremental value learning in older adults are associated with the reduced transfer of information between the hippocampus and mPFC, rather than the precision of the information in the hippocampus itself.

## Introduction

One major focus of decision neuroscience has been to understand the neural mechanisms underlying reinforcement learning (RL), our ability to estimate the value of choice options through trial-and-error updating (Niv, 2009; Sutton & Barto, 1998). Importantly, RL posits that the main driver of learning is not the outcome of each choice, but rather the difference between the realized and expected rewards, known as reward prediction error (RPE). Neural activity that tracks with RPE has been identified in the brain both at the single neuron and population level and is thought to reflect dopaminergic signaling in regions of the midbrain (Niv et al., 2012; O’Doherty et al., 2015; Schönberg et al., 2007; Schultz et al., 1997). Given the ubiquity of RL-like processes in our daily lives – from social interactions to choosing a restaurant – understanding how they evolve throughout the lifespan is of crucial importance. Here, we measure how RL differs as a function of age in a large sample and provide a systematic analysis of which aspects of value learning differ in older adults, both neurally and computationally.

In the past, studies that measured the effects of healthy aging on reinforcement learning have yielded mixed results. Several early studies claimed that older adults performed less well on RL tasks due to decreased activity in the striatum during learning (Chowdhury et al., 2013; Samanez-Larkin et al., 2014). This dampened prediction error signaling was thought to reflect decreases in dopamine availability with age (Volkow et al., 1996). More recent studies, however, have found that older adults show no deficits in RL performance, accompanied by intact value signaling in the brain (Daniel et al., 2020; Lighthall et al., 2018). Yet another study found that older adults do perform less well on simple RL tasks, but that this is primarily due to noisier value representations in prefrontal cortex during choice rather than reduced prediction error signaling (de Boer et al., 2017). More generally, the strength of the association between age and dopamine availability has been called into question, with meta-analyses highlighting that variables like receptor type and the distinction between receptor density and synthesis capacity complicate the question (Karrer et al., 2017; Seaman et al., 2019). Thus, both the extent to which healthy aging leads to impairments in incremental value learning and the extent to which these putative impairments are caused by reduced dopaminergic signaling of RPE remain open questions.

At the same time, recent work has highlighted that successful RL performance may also rely on mnemonic processes supported by the hippocampus (Biderman et al., 2020). More specifically, information from the hippocampus may serve as an input for credit assignment, allowing for a precise mapping between the choice option and its value (Ballard et al., 2019; Duncan et al., 2018; Foerde et al., 2013). Under this framework, distinct neural representations of choice options – maintained by well-characterized mechanisms like pattern separation (Bakker et al., 2008; Rolls, 2013; Yassa & Stark, 2011) – should facilitate accurate value learning. As people age, it is these kinds of processes, which are responsible for the maintenance of detailed memories, that tend to decline. For instance, work using the mnemonic similarity task (MST) has found that older adults are more likely than younger adults to mistake a perceptually similar lure for an item they have already seen (Stark et al., 2015, 2019). Neurally, older adults show altered processing in portions of the hippocampus that help maintain fine grained perceptual details, potentially leading to the “blurring” of memory that occurs with age (Yassa et al., 2011). If specific information about choice options provides an input for credit assignment during RL, and if this information becomes less precise as people age, this provides an alternative hypothesis for poorer RL performance in older adults. Rather than resulting from reduced RPE signaling, RL deficits in older adults may be related to altered processing in the hippocampus and its connections with value-responsive parts of the brain. To our knowledge, however, age-related differences in hippocampal contributions to RL have yet to be characterized.

In this paper, we set out to address these questions in a sample of older and younger adults performing an RL task in which they had to learn the value of four perceptually similar items. Of a total 251 participants, a subset of 76 performed this task while undergoing fMRI scanning, allowing us to measure the neural mechanisms underlying learning and choice. Our specific questions were threefold. First, we asked whether older adults were impaired at RL. Second, we leveraged neural data and computational modeling to identify whether any deficits that emerged were related to disruptions in learning or in choice. Third, we asked whether performance on this task was linked to the precision with which choice options are represented in the hippocampus, and to communication between the hippocampus and value-responsive circuits. Overall, we found that older adults do display subtle deficits in RL performance, which are linked to disrupted processing during choice but not during learning. Furthermore, across the lifespan, the precision with the choice options are represented in the posterior, but not anterior, hippocampus predicts accurate choice. Finally, connectivity between the posterior hippocampus and fronto-cortical circuits is reduced in older adults, highlighting a potential reduction in communication between the two brain regions. Taken together, these findings show that accurately assigning value to perceptually similar choice options may rely on an interplay between mnemonic processing in the hippocampus and fronto-cortical value signals, and that healthy aging may lead to impaired RL performance by altering this process.

## Methods

### Participants

The participants in this study are largely overlapping with those in Lempert et al. (2022) and van Geen, Cohen et al. (2024). In previous publications, we report behavioral and fMRI data from these participants while they made decisions from a single episode. Here, we turn to the second of the two tasks they completed, in a separate experimental session, which measured their ability to learn value estimates incrementally from noisy feedback (see Task section for details).

As illustrated in Figure 1B, 251 people (137 older adults and 114 younger adults) participated in this study. Of these 251, 76 participants (38 younger and 38 older adults) did the task while undergoing fMRI scanning. We had to exclude 4 of these 76 participants due to technical issues with the behavioral task, and one person for excessive head movement (see *fMRI Methods* for details on motion exclusions). In total, this resulted in a sample of 71 fMRI participants (37 younger and 34 older adults. Among the behavioral-only group (N = 175), we collected data from 97 of these participants in-person before the start of the Covid-19 pandemic, after which we switched to collecting data remotely. To approximate the experience of in-person data collection, a member of our research team was on a video call with each participant while they completed the task; in total, we collected data from 78 participants via video call. We found no systematic differences in choice accuracy between the in-lab versus remote groups (β_Method_ = 0.006, p = 0.82; β_Method*Age_ = 0.005, p = 0.89), and no systematic differences between participants who completed the study inside versus outside the scanner (β_Scan_ = -0.03, p = 0.14; β_Scan*Age_ = 0.005, p = 0.88). The demographic characteristics of each participant group are described in Table 1.

**Table 1.** Demographic characteristics of the two participant groups – those who participated only in the behavioral version of the task and those who did so while undergoing fMRI scanning.

| Age Group | Age Range | Age Details | Sex |
| --- | --- | --- | --- |
| Behavior only | Younger adults | 18 to 34 (mean: 22.3) | 54 F, 22 M |
| Behavior only | Older adults | 61 to 92 (mean: 71.3) | 60 F, 39 M |
| Behavior + fMRI | Younger adults | 21 to 37 (mean: 26.4) | 23 F, 15 M |
| Behavior + fMRI | Older adults | 61 to 86 (mean: 72.6) | 21 F, 17 M |

**Figure 1.**
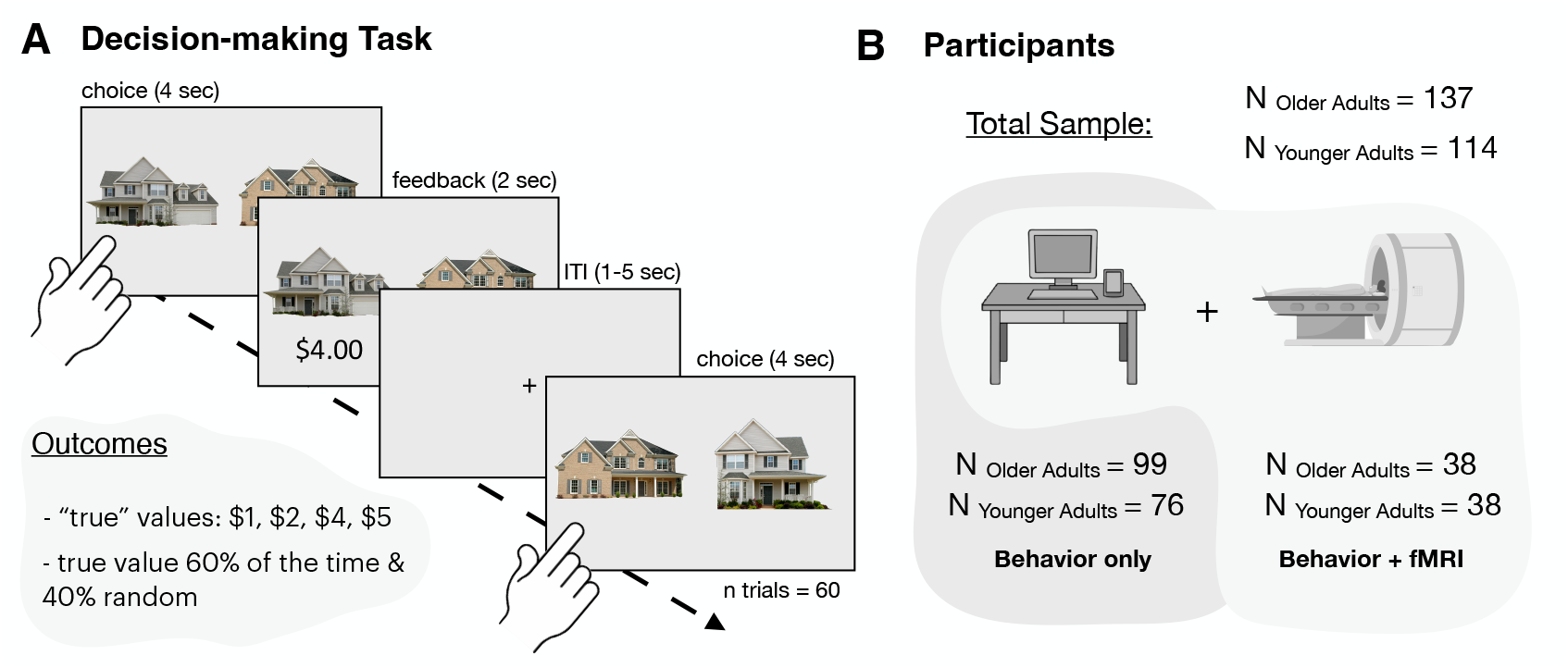
**(A)** Schematic of the reinforcement learning (RL) task. Participants have 4 seconds to choose between two of four possible houses and then receive feedback on the outcome of their choice. Each house is assigned a different value ranging from $1 to $4, which participants receive 60% of the time. On 40% of trials, the outcome associated with each house is randomly generated, ensuring the value must be incrementally learned. **(B)** Diagram of our participant sample. All participants completed the behavioral task, but only a subset underwent fMRI scanning while doing so.

We also collected behavioral data from a separate sample of middle-aged adults, whose ages spanned from 35 to 60 years old (mean age = 45 ± 7.3; 53 F, 24 M). Performance in this sample was in between the younger and older adults’ and is illustrated in Supplementary Figure 1. Because we did not collect fMRI data from middle-aged adults, we focus the results of the paper on comparisons between younger and older adults. All of the older adults in our sample were recruited from the Clinical Core of the University of Pennsylvania’s Alzheimer’s Disease Research Center (ADRC). As part of their involvement in the center, participants undergo annual psychometric testing as defined by the Uniform Data Set 3.0 (UDS) and medical/neurological examination. Older adults who participated in this study were all designated as cognitively normal based on this assessment and consensus conference determination. The young adult sample was recruited through the Penn MindCORE online participant recruitment system as well as through Facebook ads. All participants provided informed consent, and the procedures were approved by the University of Pennsylvania’s Institutional Review Board.

### Reinforcement Learning Task

In our reinforcement learning (RL) task, participants had to incrementally learn the value of two sets of four stimuli: four houses and four faces. In previous work, we have identified social biases that differentially affect choices across the age range (Lempert et al., 2022; van Geen, Cohen et al., 2024). Given these social biases with face stimuli, the focus of this paper is on age-dependent differences in RL in the non-social domain. In the absence of social biases, we hoped to better understand how purely mnemonic processes contribute to choice behavior and differ with age.

On each trial, participants were given 4 seconds to make a choice between two houses (Figure 1A). Once they indicated their selection with a key press, they received feedback about how much the house was worth. Participants were told that their goal was to maximize the amount of money they received from each choice. To do so, they had to estimate the value of each house and choose the one that was worth more. The task included four houses in total and outcomes were generated as follows: each house was assigned a ‘true’ value of either $1, $2, $4, or $5. Then, in the RL task, houses were worth that amount 60% of the time, while outcomes were randomly selected from the three other values on 40% of trials. We chose 60% as the probability of receiving the true outcome, as this yielded the greatest variation in performance in pilot studies. In total, participants completed 120 trials of this task, divided into three separate blocks of 40 trials each. In between trials, participants saw a fixation cross for a randomly determined amount of time between 1 and 5 seconds. At the end of the task, participants received the payout from one trial selected at random, which served as an additional bonus to incentivize accurate decision-making.

### Behavioral Data Analysis

All behavioral analyses were conducted in R (version 4.2.3).

To examine age-related effects on overall RL performance, we computed the average proportion of correct choices across groups for each learning block (Figure 2A). We coded a choice as correct if participants chose the higher-value house, as defined by its ‘true’ value. To test whether performance accuracy was above chance by the end of learning, we conducted a one-sample *t*-test comparing the proportion of correct choices for each participant on the last block to chance (corresponding to 0.5, since there are two possible choices). We also ran an independent samples *t*-test on participants’ accuracy scores on the last block to assess whether age group significantly modulated task performance. Finally, we used the “lme4” package in R (Bates et al., 2013) to predict trial-wise accuracy as a function of block number, age, and the interaction between age and block number. In this analysis, we also included a random intercept for each participant to account for individual differences in baseline performance.

**Figure 2.**
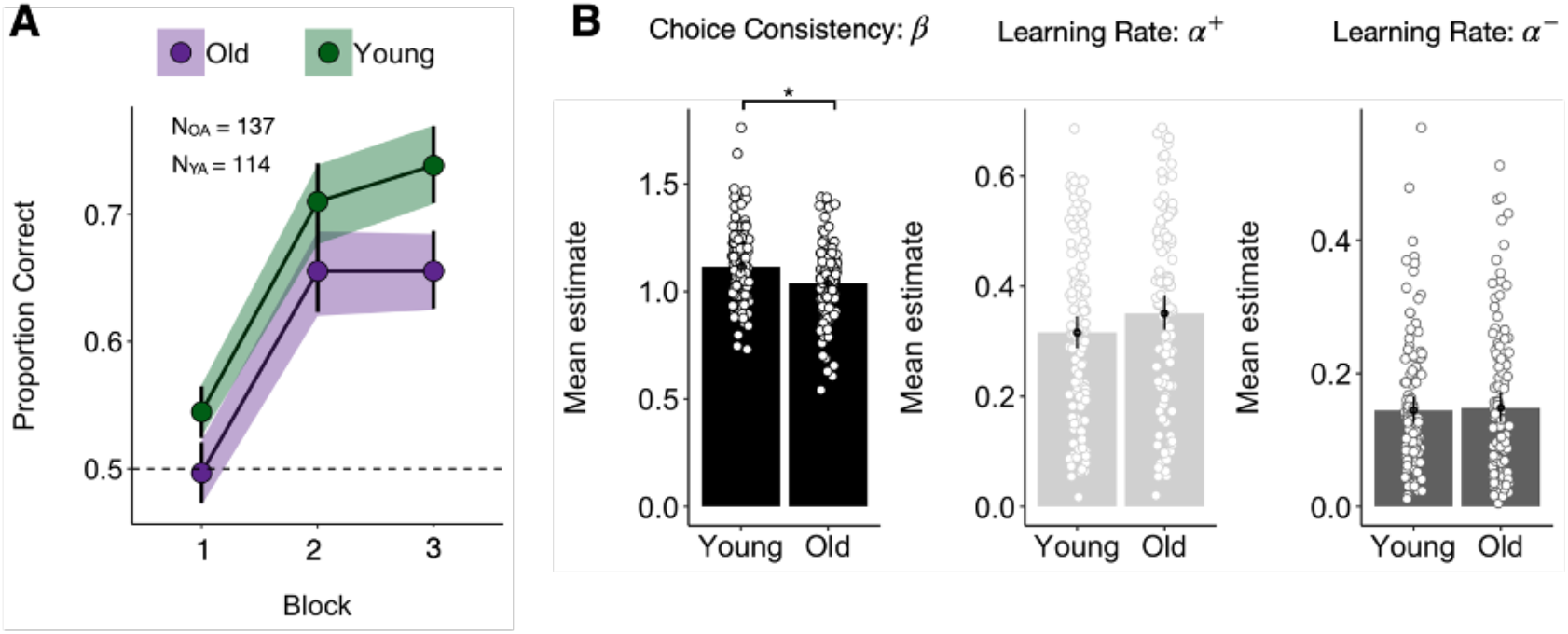
**(A)** Despite similar learning trajectories, older adults make fewer correct choices at the end of learning compared to younger adults. **(B)** This subtle age-related impairment can be captured by decreased choice consistency in a standard Q-learning model. In this same model, there are no age differences in learning rates from both positive and negative reward prediction errors (α^+^ and α^-^, respectively).

### RL Model

#### Model Description

To assess whether there were systematic age-related differences in participants’ learning rates or choice consistency, we fit a Q-learning model to the data following the hierarchical Bayesian approach described in van Geen & Gerraty (2021). Each house was assigned a separate Q-value, which was initialized at 3 (midpoint between all possible ‘true’ values). The Q-value of the chosen house was then updated after each trial as the sum of the predicted value of the chosen option 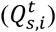 and a reward prediction error 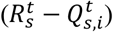, weighted by a learning rate. In the best fitting version of this model (see Model Comparison), each participant was assigned two different learning rates: one to update the Q-value in response to a positive prediction error 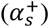, and one to update the Q-value in response to a negative prediction error 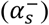. More formally, if *s* refers to subject, *t* to trial, and *i* to stimulus identity, this update corresponds to:

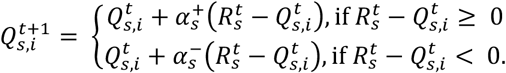

We then modeled the likelihood of choosing one stimulus or the other as a Bernoulli distribution governed by parameter *θ*. *θ* corresponds to a SoftMax transformation of the Q-values, weighted by a subject-specific β. Because there were only two choice options on each trial, the SoftMax simplifies to a logistic transform. Higher values of β correspond to a greater sensitivity to the difference in value between the two options, leading to a greater tendency towards choosing the higher-value image. We will refer to this parameter as choice consistency in the following sections. Formally, characterizing choice as either 0 (choose stimulus on the right) or 1 (choose stimulus on the left), we can model the choice probability as:

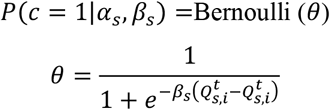

This represents the likelihood of an observed choice according to the Q-learning model, given a set of parameter values.

To extract participant-level parameter estimates, we fit the model hierarchically using Hamiltonian Markov Chain Monte Carlo (MCMC) in Stan (Carpenter et al., 2017) with 4 chains and 4000 iterations (including 2000 as burn-in). This procedure yielded posterior distributions across all 4000 samples for each parameter for each participant. We then extracted point estimates from these distributions by calculating the mean over the model-derived posteriors. To test for age differences, we conducted independent sample *t*-tests on the estimates of learning rate and choice consistency across age groups. Because the model was fit hierarchically, we simultaneously estimated group-level and participant-level parameters and used empirical priors from the group-level estimates for regularization (this method is described in detail in van Geen and Gerraty, 2021). We included the data from both younger and older adults in the same model, meaning that we did not make any assumptions about participants belonging to two different groups. This approach improves model performance because it maximizes the amount of data available for model fitting while also helping to ensure the robustness of any age differences we detect.

#### Model Comparison

The version of the model we report in the Results section contains three free parameters: an inverse temperature or choice consistency parameter that reflects how strongly Q-value guides choice, and two separate learning rates, one that updates value estimates in response to positive prediction errors and another that does so in response to negative prediction errors. We confirmed that this model outperforms a version with only one learning rate by estimating out-of-sample predictive accuracy using WAIC. WAIC provides an estimate of out-of-sample deviance, penalized by the number of parameters. We found that WAIC is larger for the version of the model with only one learning rate, suggesting that it describes the data less well than the two-learning-rate version (difference in WAIC (2 LR – 1LR) = 22181.3 - 22699.68 = -518.38).

### fMRI Methods

#### Data Acquisition and Preprocessing

Magnetic resonance images were acquired with a 3T Siemens Magnetom Prisma scanner and a 64-channel head coil. A three-dimensional, high-resolution structural image was acquired using a T1-weighted, magnetization-prepared, rapid-acquisition gradient-echo pulse sequence (voxel size=0.8×0.8×0.8mm; matrix size=241×286; 241 axial slices; repetition time=3000 ms; echo time= 30 ms). B0 field maps were acquired using a gradient echo sequence (voxel size=2×2×3 mm; matrix size=96×96; 72 slices; repetition time=1270 ms; echo time 7.46 ms; flip angle=60°). While participants completed the task, functional images were acquired using a T2*-weighted gradient echo-planar imaging pulse sequence (voxel size=2×2×2 mm; interslice gap=0.15 mm; matrix size=98×98; 72 oblique axial slices; repetition time=1500 msec; echo time= 30 ms). Slices were angled +30° with respect to the anterior commissure–posterior commissure line to reduce signal dropout in the OFC (Weiskopf et al., 2006).

We preprocessed the data using *fMRIPrep* 20.2.2 (Esteban et al., 2019), which is based on *Nipype* 1.6.1 (Gorgolewski et al., 2011). All BOLD runs were motion corrected, slice-time corrected, b0-map unwarped, registered, and resampled to a Montreal Neurological Institute (MNI) 2-mm template. After preprocessing, we performed spatial smoothing in FSL using a Gaussian kernel with a full-width half maximum (FWHM) of 5mm.

#### fMRI Analyses

##### Univariate

We conducted all univariate fMRI analyses in FSL, using generalized linear models (GLMs) to measure changes in neural activity in response to our conditions of interest. For each model, we also included confound regressors to account for motion-related artifacts. These consisted of 6 realignment parameters (three translational and three rotational), as well as the temporal derivative, quadratic terms, and the quadratic of the temporal derivative of each. Additionally, we performed spike regression by creating as many unit impulse functions as there were volumes with framewise displacement > 0.9 mm for each participant. Each of these regressors had a value of 1 at the timepoint where framewise displacement was larger than 0.9 mm, and a value of 0 elsewhere (Ciric et al., 2017). The number of spike-regressors varied substantially across participants, ranging from 0 to 74 with a median of 2. Given this wide range, we additionally excluded runs altogether if more than 10% of TRs had a framewise displacement of > 0.9mm (equivalent to any run with more than 27 spike regressors), and we excluded participants altogether if the majority of their runs (2 or more) fit this criterion. In practice, this resulted in the full exclusion of 1 participant and the exclusion of 1 run for an additional 3 people. Group-level maps were computed using a FLAME1 + FLAME2 mixed effects model, thresholded at *p* = 0.001, and corrected for multiple comparisons using parametric cluster-based correction.

We used a recently developed python-based pipeline (https://github.com/alicexue/fmri-pipeline) to help automate the analyses in FSL. As described in the results section, we modeled neural activity separately for the 4-second choice phase and the 2-second feedback phase. For the results reported in Figure 3, we included the following regressors of interest in our GLM: 1) choice phase, 2) parametric modulator during the choice phase for the Q-value of the chosen option, 3) feedback phase, 4) parametric modulator during the feedback phase for reward prediction error. To make sure that we modeled neural activation comprehensively, we also included the same set of regressors for the trials on which participants chose between two faces, although these trials are not the focus of this paper. These trials were intermixed with house trials in each run.

**Figure 3.**
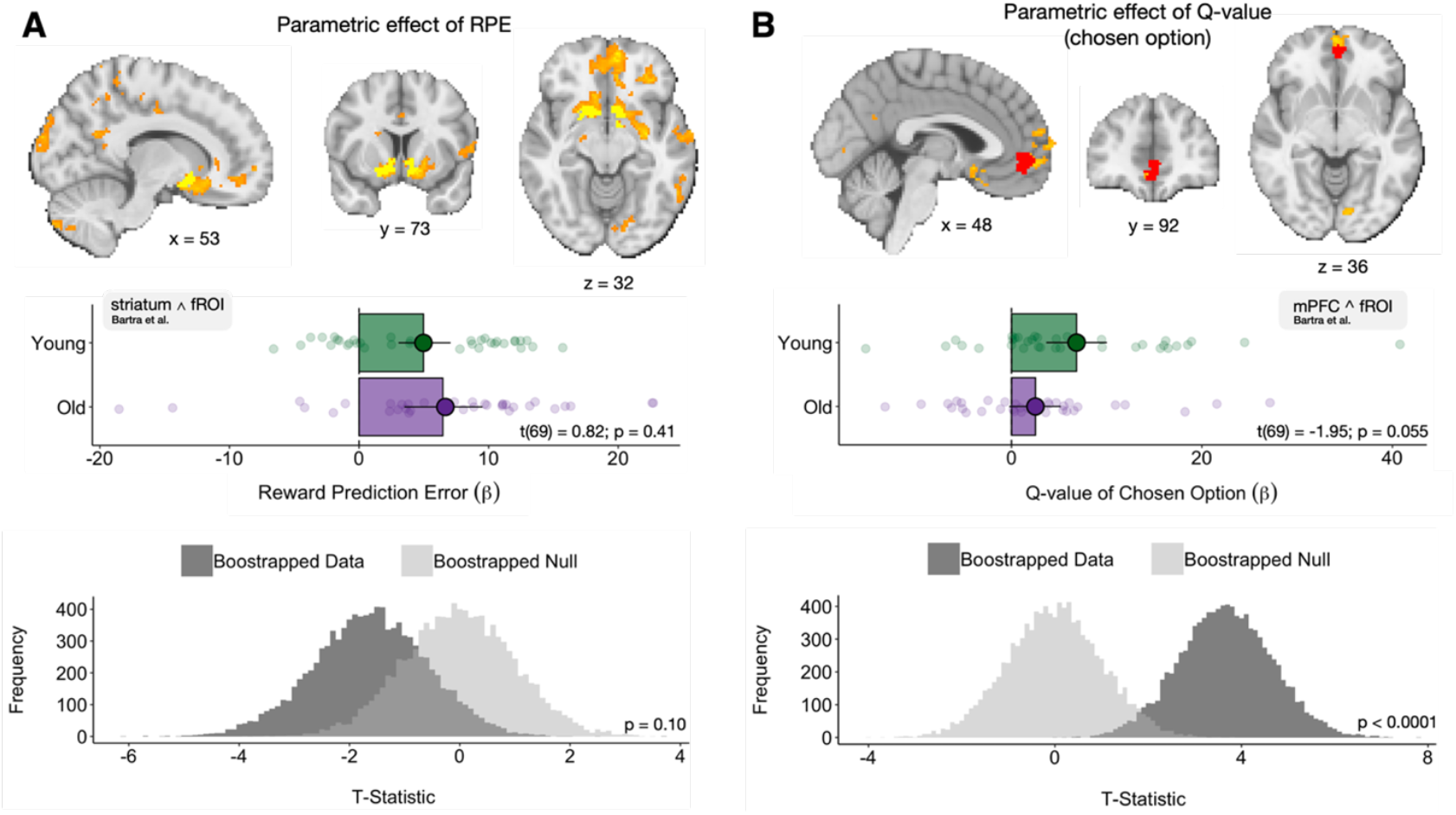
**(A)** Neural activity in value-responsive brain regions including the ventral striatum tracks with reward prediction error (RPE) across age groups. There are no significant age differences in the magnitude of this activation in a fROI defined as the intersection between the univariate activation and outcome-sensitive regions of the striatum identified in a large meta-analysis (Bartra et al., 2013). This is true in the actual fMRI sample and a bootstrapped group of participants whose sample size matches that of the behavioral data (see Methods for more details on this approach). **(B)** Neural activity in the mPFC tracks with the Q-value of the chosen option across the full sample of participants. However, in a fROI of the mPFC, our bootstrapping analysis suggests that activation may be stronger in the younger than the older adult group. In both **A** and **B**, the fROIs from which we extracted participant-specific coefficients are overlaid in yellow and red, respectively.

We then assessed whether there were any effects of age on the strength of neural signaling by combining *a priori* regions of interest (namely, the striatum and mPFC) with the results from the univariate analyses. In the case of prediction error signaling, since neural activity tracked with prediction error in a large cluster that extended beyond the striatum, we created a functional ROI by intersecting the univariate activation with value-responsive regions of the striatum identified in Bartra et al., 2013 (Figure 9 of Bartra et al.: 5-way conjunction analysis designed to detect brain regions that track subjective value). In the case of Q-value signaling, we followed a similar approach and generated the fROI by intersecting the univariate activation with regions of the mPFC identified in Figure 9 of Bartra et al. (2013). Within each fROI, we then used *fslmeants* to extract subject-specific betas that reflect the magnitude of neural activation in response to the variable of interest. We then performed an independent-samples *t*-test on the extracted coefficients in order to test for age differences. Given the difference in size between our behavioral and fMRI samples, we further evaluated age differences following the bootstrapping procedure described below.

#### Bootstrapping

Our goal in this paper was to integrate the behavioral and fMRI findings; however, a challenge in doing so is that the behavioral sample (N = 251) is much larger than the fMRI sample (N = 72). This difference in sample size makes it difficult to interpret age differences in the fMRI sample when they are not large enough to reach statistical significance. For example, the age differences we observed in behavior were only robust when we considered the entire sample, and not when we restrict this analysis to the fMRI group (see Supplementary Figure 1 for fMRI-only behavioral results). Is this null result likely due to a lack of statistical power, or should we interpret it as evidence against age differences in the fMRI sample? To address this question, we devised a statistical test based on a bootstrapping procedure to assess whether age differences might have been reliably detectable had the fMRI sample been of the same size as the behavioral one.

To do this, we generated 10,000 hypothetical datasets by sampling data points (with replacement) until the simulated fMRI sample size was equivalent to our behavioral sample (137 older adults and 114 younger adults). For each generated dataset, we then performed an independent sample *t*-test comparing the magnitude of neural activation across the two age groups. This procedure generated a distribution of 10,000 t-statistics, approximating the strength of the neural differences in a would-be sample of the same size as our behavioral sample. We then conducted a permutation test using the same sampling procedure but shuffling the age labels across subjects. We compared the null distribution to the bootstrapped distribution and computed a *p*-value as the proportion of permuted t-statistics that were equal to or exceeded the absolute value of the mean t-statistic of the bootstrapped distribution. This test estimates the likelihood that the age differences we observed in the fMRI sample would have reached statistical significance, were we to have an fMRI sample the size of our behavioral sample.

#### Representational Similarity Analysis

The process by which we identified stimulus-specific patterns of neural activity for each participant is illustrated in Figure 4A. First, we used *nibeta-series* to extract beta series from the functional MRI data, using the same set of confound regressors as in the univariate analyses. This procedure isolated patterns of neural activity during every trial of every run. Then, because the task design is such that two stimuli are always on the screen at the same time for each trial, we used a regression-based approach to estimate the neural response to each of the four possible houses independently (step 2 in Figure 4A). We created a stimulus design matrix for each scan run that was composed of as many rows are there were trials (forty) and as many columns as there were stimuli (four). We populated the stimulus design matrix with ones and zeros, with ones indicating the presence of that stimulus on that particular trial. Using Ordinary Least Squares (OLS), we then regressed the neural activity onto the stimulus design matrix, allowing us to derive an activation coefficient for each voxel in response to each stimulus. Importantly, this only extracts stimulus-specific patterns of neural activity at the run level, rather than on each trial. Next, we correlated the run-level coefficients for each of the same stimuli on different runs (r_same_) and compared the strength of that correlation to the correlation between different stimuli on different runs (r_diff_). The difference between these two values (r_same_-r_diff_) is our main differentiation metric and captures the extent to which each person’s brain represents different stimuli differently, relative to a within-participant baseline.

**Figure 4.**
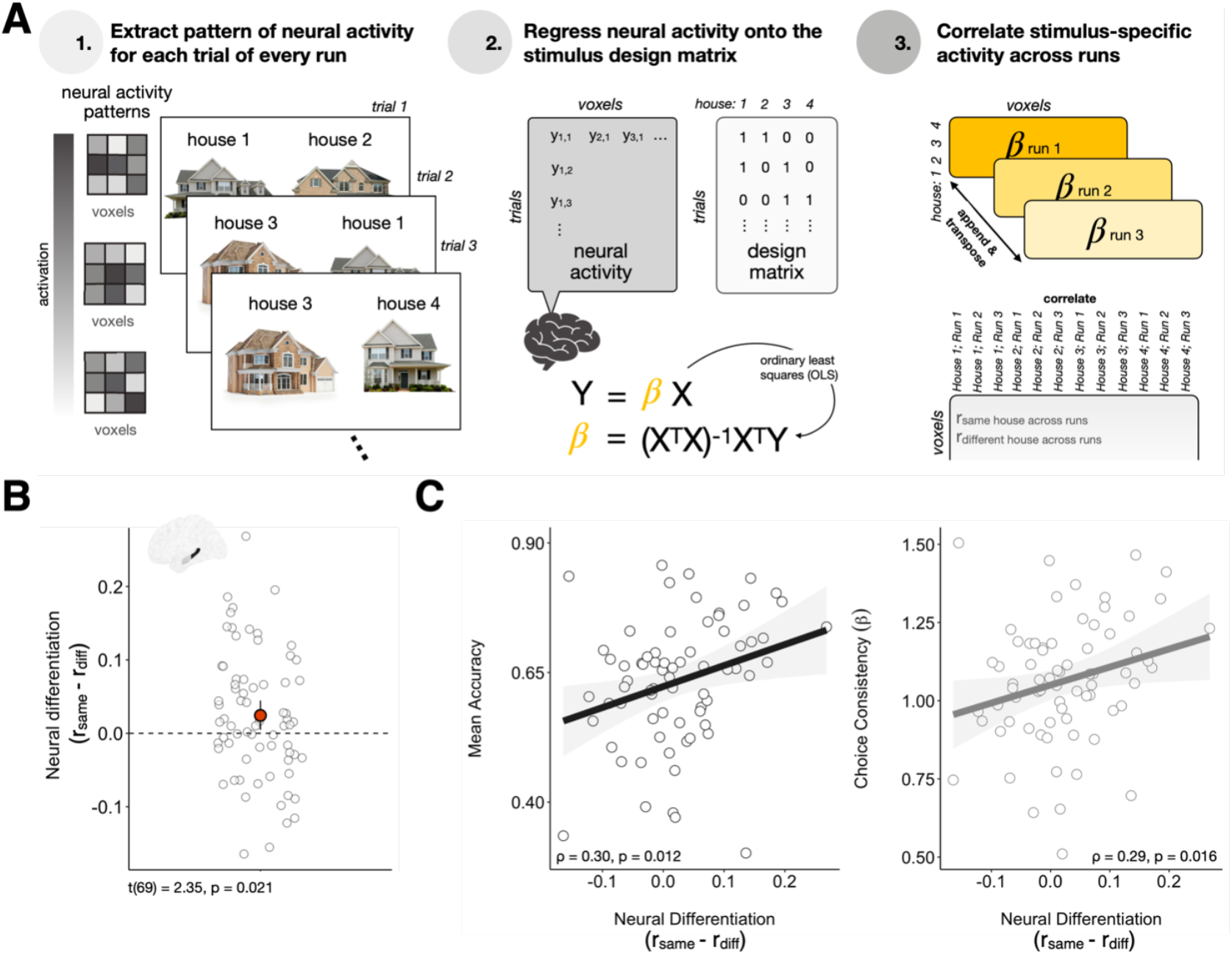
**(A)** Overview of the method that allows us to identify stimulus-specific patterns of neural activity. First, we extract patterns of neural activity for each trial for every run. Then, in order to identify stimulus-specific activation, we regress neural activity for each run onto a stimulus design matrix using Ordinary Least Squares (OLS). Finally, we correlate the beta coefficients for each stimulus across runs and compare the strength of same-stimulus versus different-stimulus correlations. **(B)** Distribution of average neural differentiation across our sample. In the posterior hippocampus, patterns of neural activity at the group level are more similar when the same stimulus is repeated across runs than when different stimuli are repeated (mean differentiation > 0). **(C)** *Left*. Stimulus differentiation in the posterior hippocampus predicts choice accuracy across participants. *Right*. Stimulus differentiation in the posterior hippocampus also predicts choice consistency, as captured by the β parameter in the Q-learning model

In this analysis, we were specifically interested in patterns of neural activity in the hippocampus, and its anterior and posterior portions independently. To localize neural activity to these regions of interest (ROIs) we multiplied the beta series with masks that contained zeros in all but the relevant brain areas. We used anterior and posterior hippocampal ROIs that were defined from a probabilistic atlas of the medial temporal lobe (Hindy & Turk-Browne, 2016).

Finally, because the similarity metric described above crucially relies on computing correlations in activity patterns across runs, we did not apply the 10% motion threshold that led to the exclusion of entire runs in the univariate analyses – rather, we included all runs for all participants, excluding the one participant with excessive head motion on every single run. However, we verified that all effects hold if we additionally exclude the three participants who each had excessive head motion on one of the runs.

#### Functional Connectivity

To measure age effects on functional connectivity, we conducted a seed-based functional connectivity analysis in FSL. Based on our multivariate findings, we were specifically interested in understanding whether the posterior hippocampus was differentially connected to the rest of the brain in younger versus older adults. Thus, the GLM in this analysis consisted of a single regressor of interest, the timeseries from the posterior hippocampus, as well as the same regressors of no-interest that we used in other analyses to account for motion. We then tested for whole-brain age differences using a FLAME1 + FLAME2 mixed effect model, thresholded at *z* = 3.1, and corrected for multiple comparisons using parametric cluster-based correction.

Given our univariate results, we were specifically interested in the magnitude of this effect in the mPFC. To quantify this, we used *fslmeants* to extract subject-specific betas in an ROI we defined anatomically using the Harvard-Oxford Atlas and tested to what extent age determines this brain region’s connectivity with the posterior hippocampus. Specifically, we ran an independent-samples *t*-test on the extracted coefficients and compared the strength of connectivity between the posterior hippocampus and relevant portions of the mPFC in younger versus older adults.

## Results

### 1. Behavior

First, we verified that participants in our full behavioral sample (n = 251) learn to choose the higher-value image over time. To do so, we computed the proportion of trials in each block on which participants chose the higher-value option. For both younger and older adults, choice accuracy increases significantly as learning progresses, and is, on average, well above chance by the end of the task (t(244) = 17.1, p > 0.0001, Figure 2A). However, despite similar learning trajectories (β_Block*Age_ = 0.008; p = 0.25), older adults make fewer correct choices by the end of learning, as illustrated by lower choice accuracy in block 3 (t(244) = -3.7, p = 0.0002, Figure 2A).

To better understand the mechanisms underlying these age differences, we fit a Q-learning model to the data that characterizes task performance based on a choice consistency parameter (β or inverse temperature) and two separate learning rates: one that updates value estimates after positive reward prediction errors and another that updates value estimates after negative reward prediction errors (see Methods for more details). We included two different learning rates because this version of the model outperforms a version with only one learning rate (difference in WAIC = -518.38). From this model, we find that older adults’ reduced accuracy is captured by significantly lower choice consistency (t(222) = -3.34, p = 0.0009, Figure 2B) rather than age differences in learning rates (α^+^: t(222) = 0.21, p = 0.82; α^-^: t(222) = 1.58, p = 0.12, Figure 2B).

### 2. fMRI: Univariate

Our behavioral findings suggest that age differences in RL performance are likely related to a noisier decision process rather than impaired value updating. To further assess this claim, we turned to the neural data and tested whether there are significant age differences in neural activity during feedback or during choice. Focusing first on the 2 seconds of feedback, we found that model-derived reward prediction error (RPE) tracks with neural activity in a variety of brain regions including the striatum and mPFC (Figure 3A). In a functional Region of Interest (colored yellow in Figure 3A) identified based on the intersection between the univariate activation and the value-responsive regions of the striatum identified in Bartra et al. (2013), age differences in RPE signaling are not significant (t(69) = 0.82, p = 0.41). Bootstrapping the fMRI data to estimate the effect in a sample the size of our behavioral sample (see Methods), we found that this null result is unlikely to be due to the smaller sample size in the fMRI data (permutation test: p = 0.10). In fact, if anything, the trend is towards stronger RPE signaling in older adults. This set of findings parallels the absence of age differences in learning rate estimates derived from the Q-learning model. Both the neural and behavioral data suggest that deficits in RPE signaling during feedback are not at the root of older adults’ lower performance.

Instead, older adults may represent Q-value less strongly, leading to more stochastic decisions. Turning to the choice phase, we found that consistent with previous work (Knutson et al., 2001; Niv, 2009), neural activity in the medial prefrontal cortex (mPFC) parametrically tracks the Q-value of the chosen option (Figure 3B). To assess age differences in this effect, we again computed a fROI (in red in Figure 3B) by taking the intersection between the univariate activation and the parts of mPFC that were identified as value-sensitive in Bartra et al. (2013). Focusing on this fROI, we found that Q-value signaling is marginally stronger in the younger than the older adult group (t(69) = -1.85, p = 0.055, Figure 3B). When we used bootstrapping to estimate this effect in a sample the size of our behavioral sample, we found that this effect would likely reach statistical significance in a larger sample (permutation test: p < 0.0001). This set of fMRI findings is consistent with the notion that dampened value representations during choice may underlie noisier performance in older adults.

### 3: fMRI: Representational Similarity Analysis

So far, we have identified a possible neural correlate of older adults’ lower choice consistency: a tendency towards reduced Q-value signaling in the mPFC. It remains to be seen, however, what contributes to the strength of this Q-value representation, and whether age may affect this process. One possibility is that accurate value signaling relies on the retrieval of information about choice options from the hippocampus. If this is true, we might expect the precision with which each stimulus is represented in the hippocampus to predict value learning and consequent choice accuracy. Conversely, if these representations become blurred, participants may have more difficulty with credit assignment, leading to more stochastic choice. Given that this hypothesis relies on the maintenance of detailed and specific memories, we focus our analyses on the posterior portion of the hippocampus, which has been shown to be preferentially implicated in memory precision and pattern separation (Robin & Moscovitch, 2017; Wanjia et al., 2021; Yassa et al., 2011; Yassa & Stark, 2011).

We used Representational Similarity Analysis (RSA) to measure the precision with which each stimulus is represented in the posterior hippocampus. As described in greater detail in the Methods section and Figure 4A, we extracted stimulus-specific patterns of neural activity for each house during each run of scanning. First, we verified that these patterns were more similar for the same image presented across runs than different images presented across runs. We computed a metric of relative neural differentiation for each participant, corresponding to the difference in the average neural correlation between repeats of the same house and that of different houses (r_same_-r_diff_) – values greater than 0 are indicative of stimulus-specific differentiation. Overall, repeats of the same house were indeed represented more similarly in the posterior hippocampus than repeats of different houses (one-sample t-test compared to 0: t(69) = 2.35, p = 0.021), suggesting that this brain region maintains stimulus-specific representations.

We next asked whether stimulus differentiation in the posterior hippocampus is important for successful credit assignment and consequently accurate choice. In line with this hypothesis, we found a positive correlation between stimulus differentiation in the posterior hippocampus and choice performance across participants (ρ = 0.30, p = 0.012, Figure 4C). We also found a positive relationship between stimulus differentiation and each participants’ choice consistency parameter from the Q-learning model (ρ = 0.29, p = 0.016, Figure 4C). These relationships do not hold in the anterior portion of the hippocampus, which we would expect to be less important for maintaining stimulus-specific representations (ρ = 0.08, p = 0.50 and ρ = 0.06, p = 0.57 respectively). These findings suggest that stimulus representations in the posterior hippocampus could provide an input for value learning and facilitate Q-value computations.

Given the results above, we might expect that older adults represent different stimuli less distinctly, leading to more imprecise valuation and lower performance. To test this hypothesis, we asked whether there were any age differences in stimulus differentiation in the posterior hippocampus, or in its relationship to choice. Interestingly, the relationship between stimulus differentiation and choice performance holds across the age range, with no significant interaction between age and the strength of the effect (β_Diff*Age_ = 0.25, p = 0.43). Furthermore, there was no main effect of age on relative stimulus differentiation (t(68) = -0.1.19, p = 0.24). When we used bootstrapping to estimate age differences in stimulus differentiation in a sample the size of our behavioral sample, we found that a larger sample would possibly reveal stronger, rather than weaker, neural differentiation in the posterior hippocampus of older adults (permutation test: p = 0.025). Thus, the extent to which stimuli are distinctly represented in the posterior hippocampus does not seem to differ with age.

### 4: fMRI: Functional Connectivity

Although stimulus differentiation appears intact in older adults, we see possible dampening in the extent to which the mPFC activity tracks Q-value in older adults. Mechanistically, these findings suggest that age may be linked to a disruption in how stimulus information from the posterior hippocampus impacts an evaluative process performed in the prefrontal cortex.

As a test of this hypothesis, we conducted a connectivity analysis looking at where neural activity is more functionally coupled with the posterior hippocampus in younger than older adults. Given our hypothesis, we were particularly interested in measuring connectivity between the posterior hippocampus and the mPFC, as defined by an anatomical mask. Figure 5A shows that, as predicted, there is greater connectivity between the posterior hippocampus and the mPFC in the younger than older adult group (t(69) = -2.46, p = 0.017). One young adult participant was a clear outlier in this analysis, identified by both a modified Z-score exceeding 3.5 and significance on Grubbs’ test with p < 0.001 (red dot in Figure 5). To ensure that this outlier was not driving the significant age difference, we verified that the effect was robust to excluding them (t(68) = -2.3, p = 0.027). Finally, if the transfer of information between the posterior hippocampus and mPFC supports accurate value learning, we might expect the connectivity between the two brain regions to be correlated with choice accuracy and consistency. Supporting this hypothesis, we find that across the full sample, individual differences in functional connectivity between the two brain regions predict marginally more accurate and significantly more consistent choice (ρ = 0.19, p = 0.097 and ρ = 0.31, p = 0.0091 for accuracy and choice consistency respectively, Figure 5B). Taken together, these results highlight a potential disruption in the transfer of information between posterior hippocampus and mPFC in older adults, with downstream consequences for choice accuracy and consistency.

**Figure 5.**
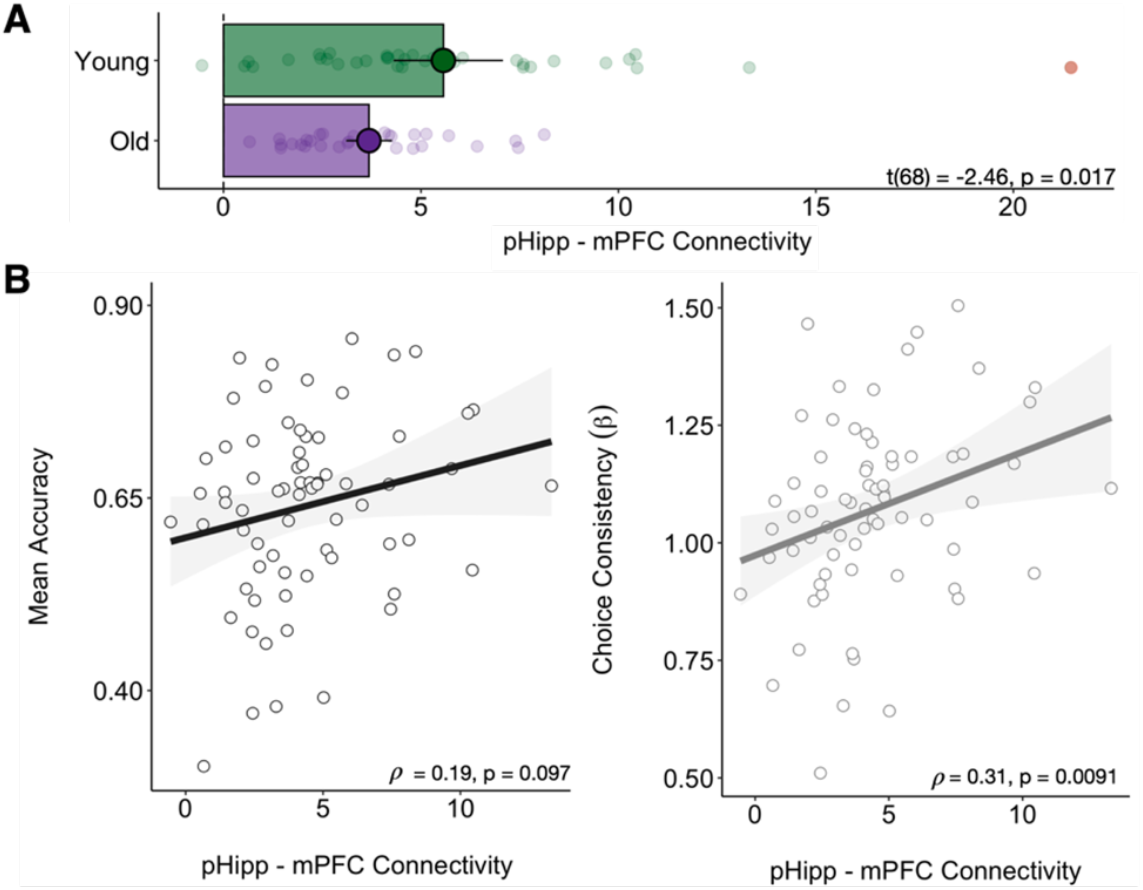
**(A)** Age differences in functional connectivity between posterior hippocampus and mPFC across the experimental task. There is greater functional connectivity between the two relevant brain regions in younger than older adults. **(B)** Across individuals, connectivity between posterior hippocampus and mPFC tends to predict performance accuracy and choice consistency (the latter is derived from the computational model). For visualization purposes, the outlier identified in red in Panel A is excluded from these plots, but it is included in the statistical test.

## Discussion

Here, we aimed to address several questions: (1) Do older adults perform less well than younger adults on a reinforcement learning task? (2) Can the origins of any performance differences be traced back to altered processing during feedback or during choice? (3) Could hippocampal contributions to RL help explain individual differences in task performance, both across individuals within the full sample and between the two age groups? Our data suggest that older adults perform subtly but significantly less well in an RL task, and that these deficits emerge due to reduced value signaling at choice rather than reduced prediction error signaling at feedback. Furthermore, precise representations of the perceptually similar choice options in the posterior hippocampus positively predict performance across the age range. The stimulus-specificity of these representations is not, contrary to what we hypothesized, lower in older adults – however, functional connectivity between the hippocampus and mPFC is, suggesting that, in this experimental context, it may be the transfer of information rather than the information itself that is disrupted in older adults.

Our finding that hippocampal representations of choice options predict performance complements and expands traditional views of RL, which tend to view it primarily as a form of procedural learning that relies on the basal ganglia (Gerraty et al., 2018; O’Doherty et al., 2015; Schönberg et al., 2007; Squire, 1992). While we do see strong RPE signaling in the striatum, as expected, we also see evidence that stimulus information stored in the hippocampus may play a part in credit assignment. This perspective is in line with more recent findings that highlight the hippocampus’s role in incremental value learning. On one hand, extensive evidence links the hippocampus to reward learning in complex environments – for instance when multiple steps lead to a reward and successful choice requires an understanding of high-level state transitions (Doll et al., 2012; Liu et al., 2021; Russek et al., 2017; Simon & Daw, 2011; Vikbladh et al., 2019; Wimmer & Shohamy, 2012). Beyond the involvement of the hippocampus in this sort of “model-based” planning, however, it may also facilitate simpler stimulus-outcome associations in some task contexts. For instance, activity in the hippocampus tracks with learning from trial-by-trial feedback in adolescents (Davidow et al., 2016), and during configural or temporally extended RL in adults (Ballard et al., 2019; Duncan et al., 2018; Foerde et al., 2013). In these tasks, value computation often relies on the explicit recruitment of mnemonic processes – for instance, to remember the feedback that was associated with a choice after a delay of several seconds. Similarly, in our study, participants need to keep track of more perceptually similar stimuli than are presented on the screen at any given time. Thus, hippocampal contributions to “model-free” RL may emerge specifically when credit assignment relies on forming holistic representations of choice options that need to be preserved over time (but see Palombo et al. (2019) for a different perspective in amnesiacs).

That the hippocampus would be involved in both configural learning and in maintaining distinct, stimulus-specific representations as in our task points to its dual and sometimes paradoxical function. On one hand, the hippocampus allows for associations, patterns, and configurations to be learned and remembered, combining features across experienced events (Brasted et al., 2003; Gilboa & Marlatte, 2017; Schapiro et al., 2012; Schlichting et al., 2017). On the other hand, the hippocampus also encodes episodic memories, which are inherently unique, and maintains separate records of even very similar experiences (Chanales et al., 2017; Korkki et al., 2021; Scoville & Milner, 1957; Tulving & Markowitsch, 1998). To unite these opposing functions – often referred to as pattern completion and pattern separation – researchers have identified a granularity gradient the spans the long axis of the hippocampus (Robin & Moscovitch, 2017; Strange et al., 2014). According to this theory, the anterior hippocampus supports abstract, generalized processing and is responsible for high-level tasks like category learning. Conversely, the posterior hippocampus is thought to encode detailed and precise memories, keeping track of individual items at the exemplar level. This anterior-posterior distinction may arise because different hippocampal subfields are preferentially represented in specific sections of the hippocampus. Indeed, the dentate gyrus (DG) and CA3 take up most of the posterior hippocampus and are the central nodes of a pathway that enables pattern separation (Bakker et al., 2008; Leutgeb et al., 2007; Rolls, 2013; Schapiro et al., 2017; Sučević & Schapiro, 2022; Yassa & Stark, 2011). For this reason, our finding that stimulus differentiation in the posterior (but not anterior) hippocampus predicts accurate choice is consistent with both theoretical and empirical evidence. Finally, although most studies of pattern separation do not ask people to make choices beyond a memory test, some have found that stimulus differentiation tracks with “success”, broadly defined. Wanjia et al. (2021), for instance, found that accurately remembering stimulus-stimulus associations relies on neural dissimilarity in the CA3/DG, and a sudden increase in this dissimilarity tracks with the resolution of memory interference. In our sample, we were not confident that the spatial resolution of our fMRI scans would allow for subfield segmentation, but we hope to examine representations in specific hippocampal subfields in future work.

Despite providing evidence for hippocampal involvement in credit assignment, our findings do not negate that RL relies on partly specialized, extra-hippocampal systems. Since the hippocampus is far from the only brain region that undergoes changes as people age, altered functioning in value-responsive parts of the brain could also explain the performance differences we observe. However, we find no evidence for an attenuation in striatal signals of prediction error in older adults. This parallels the most recent research on the topic, which fails to identify any disruption in the neural correlates of RPE (Daniel et al., 2020; de Boer et al., 2017; Lighthall et al., 2018; Seaman et al., 2023). If anything, we and others (Daniel et al., 2020) see numerically stronger prediction error signals in the ventral striatum of older adults. In light of evidence that feedback-related activity in the striatum is stronger in high-effort tasks (Dobryakova et al., 2017), one reason for this may be that older adults perceive the task as more difficult. Even without making assumptions about the task’s perceived difficulty, numerically stronger RPE signaling follows from assuming older adults form noisier value estimates and as a consequence, experience more prediction errors. Further corroborating this idea is the fact that in our sample, we see a negative relationship between RPE signaling in the striatum during feedback and Q-value representation at choice (r = -0.29, p = 0.021). That said, several older papers did find reduced or incomplete prediction error signaling when older adults perform a range of probabilistic value-learning tasks (Chowdhury et al., 2013; Eppinger et al., 2013; Samanez-Larkin et al., 2014; Schott et al., 2007). However, these studies either had small sample sizes (N < 14 in each age group) or did not include a younger adult comparison group. Furthermore, none report systematic age differences in striatal RPE signaling from whole-brain or region-of-interest analyses that use model-derived prediction error as a parametric regressor. Conversely, the more recent studies finding no age differences in RPE signaling (Daniel et al., 2020; de Boer et al., 2017; Lighthall et al., 2018) all have sample sizes of at least 25 participants in each group and identify neural signals of RPE using the same approach we do: deriving its magnitude from an RL model and using that as a parametric regressor in ROI analyses that test for age effects. Thus, we believe that the preponderance of evidence favors the conclusion that there are no reliable age differences in prediction error signaling.

Our finding that neural differentiation in the posterior hippocampus does not differ with age contributes to a similarly lively debate. On one hand, the phenomenon of age-related neural dedifferentiation (Koen & Rugg, 2019) posits that as people get older, the amount of noise present in their neural activity increases, leading neurons to become less specialized and patterns of activation to become less distinct. It is important to note, however, that most of the evidence in favor of dedifferentiation focuses on category-selective regions of the cerebral cortex. For instance, neural activity in the fusiform face area (FFA) of older adults does not discriminate between similar-looking faces (Goh et al., 2010). This finding is supported by studies that, like ours, measure similarity in patterns of neural activity as a proxy for neural differentiation, or lack thereof (Srokova et al., 2024; St-Laurent et al., 2014). Whether we should expect similar decreases in the selectivity of neural response patterns in the hippocampus, however, remains less clear. Based on behavioral findings, we have good reason to predict that this would be the case: using experimental paradigms like the Mnemonic Similarity Task (MST), which asks people to encode a large number of object images and perceptually similar lures, researchers have found a reliable decrease in memory precision in older adults (Greene & Naveh-Benjamin, 2023; Stark et al., 2015, 2019). Neurally, the DG/CA3 hippocampal subregions may also be hyperactive in older adults, specifically during trials that require pattern separation (Yassa et al., 2010). Taken together, these findings have led to a great deal of speculation about whether hippocampal activity patterns become less stimulus selective as people get older, underlying the reported ‘blurrier’ memory (Leal & Yassa, 2018). However, as far as we know, evidence in favor of this claim based on multivariate analyses of patterns of neural activity is scarce. For instance, the study that highlights increased pattern similarity in category-selective regions of cortex does not implicate the hippocampus, nor does other research that explicitly links dedifferentiation to memory performance (Koen et al., 2019). It may also be that we do not have the ability to detect decreases in hippocampal differentiation in our current task, either because there are only four stimuli, or because they are not sufficiently similar. We hope to formally assess this claim in future RL tasks that require participants to discriminate between more than four individual options of greater perceptual similarity.

Although hippocampal differentiation may not differ in older adults, our findings do suggest a disruption in how stimulus-specific information is communicated from the hippocampus in service of value-signaling in mPFC. Previous evidence looking at the functional connectivity between the hippocampus and cortex in older adults has similarly found a reduction in the extent to which activity in the posterior hippocampus specifically is coupled with mPFC activation (Grady et al., 2003; Panitz et al., 2021). In our sample, it may be that this reduced connectivity underlies the attenuation of value signaling in the mPFC of older adults. This mechanism would be in line with a view of the hippocampus’ contribution to RL as providing an input for credit assignment, as has been previously proposed (Ballard et al., 2019). Our findings, however, also raise an important follow-up question: if the hippocampus contributes to RL only insofar as it allows for the accurate transfer of value information to prefrontal regions, how come the precision with which it represents choice options predicts performance accuracy in older adults, for whom this process is disrupted? The simplest explanation is that despite the group-level decrease in connectivity, individual differences in the transfer of information relate to hippocampal precision. However, other alternatives could also explain these findings. For instance, it could be that in older adults, an alternate neural pathway, implicating the hippocampus and brain regions other than the mPFC, supports the link between stimulus representation and choice accuracy. Yet another possibility is that the hippocampus itself encodes not only the identity of choice options but also information about their respective values (Knudsen & Wallis, 2021). Further adjudicating between these alternatives is an important question for future research.

## Acknowledgements

This work was funded by grant RF1AG058065 from the National Institute on Aging.

## Supplementary Figures

**Supplementary Figure 1.**
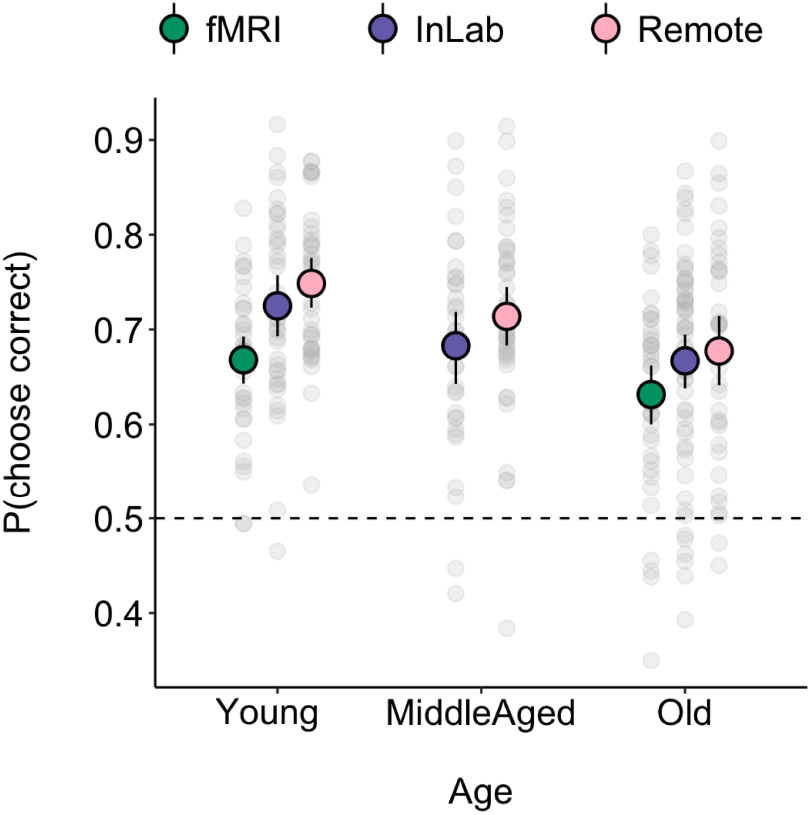
Average performance accuracy across our different participant samples. Middle aged adults performed marginally worse than younger adults (t = 1.9; p = 0.06) and not significantly differently from older adults (t = -1.25, p = 0.21), if we exclude the fMRI sample. In the fMRI sample, older adults performed only marginally worse than younger adults (t = -1.17, p = 0.084).

